# dbMTS: a comprehensive database of putative human microRNA target site SNVs and their functional predictions

**DOI:** 10.1101/554485

**Authors:** Chang Li, Michael D. Swartz, Bing Yu, Yongsheng Bai, Xiaoming Liu

## Abstract

microRNAs (miRNAs) are short non-coding RNAs that can repress the expression of protein coding messenger RNAs (mRNAs) by binding to the 3’UTR of the target. Genetic mutations such as single nucleotide variants (SNVs) in the 3’UTR of the mRNAs can disrupt this regulatory effect. In this study, we presented dbMTS, the database for miRNA target site (MTS) SNVs, which includes all potential MTS SNVs in the 3’UTR of human genome along with hundreds of functional annotations. This database can help studies easily identify putative SNVs that affect miRNA targeting and facilitate the prioritization of their functional importance. dbMTS is freely available at: https://sites.google.com/site/jpopgen/dbNSFP.

## Introduction

MicroRNAs (miRNAs) are short non-coding RNAs (∼ 22 nucleotide) that can repress the expression of target messenger RNAs (mRNAs) by binding to their 3’ untranslated regions (UTRs). It is estimated that more than 60% of human protein coding genes are under the regulation of miRNAs (Friedman, Farh, Burge, & Bartel, 2009; Lewis, Burge, & Bartel, 2005), and there is increasing evidence suggesting their wide variety of functions in developmental and physiological processes (Bartel, 2018). miRNAs always convey their repressive effect through an imperfect binding with their target mRNAs, however perfect base-pairings at the 5’ end of the miRNA (nucleotide position 2-7; also known as the seed region) is often required for a miRNA target site (MTS) to be functional (Agarwal, Bell, Nam, & Bartel, 2015). Thus, single nucleotide variants (SNVs) located within MTS especially those residing in the part of the MTS that pairs with miRNA seed regions, can undoubtedly disrupt the efficacy of miRNA targeting. This type of regulatory variants can then lead to downstream transcriptomic and proteomic changes which have been extensively reported to be associated with various diseases (Li et al., 2018; Nicoloso et al., 2010). However similar to many other regulatory SNVs, the functional importance of most MTS SNVs is still poorly understood.

To help us understand and interpret these regulatory SNVs in MTS, some databases have been established to try to link these SNVs with miRNA targetome alterations and diseases (Bhattacharya, Ziebarth, & Cui, 2014; Bruno et al., 2012; Gong et al., 2015; C. Liu et al., 2012). While some databases lack recent updates, two widely used and actively updated databases are: PolymiRTS Database 3.0 (Bhattacharya et al., 2014) and miRNASNP v2.0 (Gong et al., 2015). These two databases both used variants from dbSNP build 137 (Sherry et al., 2001) and tried to link these variants with MTS and/or possible downstream phenotype information. Although these aforementioned databases are valuable in interpreting interesting association results, they still suffer from some limitations. First, only known variants from dbSNP were included in the databases. With the fast development of whole-genome sequencing (WGS), there is a growing need in predicting the functional effect of novel SNVs. Therefore, focusing only on known variants from dbSNP is clearly insufficient. Second, their information is not comprehensive and has omitted a vast majority of recently developed functional annotations that can help interpret these MTS SNVs, e.g. CADD (Kircher, 2014), Eigen (Ionita-Laza, McCallum, Xu, & Buxbaum, 2016) etc. Such annotations utilized a wide range of functional genomic annotations such as conservation, experimentally identified functional elements and their consequences etc., and they were proven to be predictive regarding the functional consequences of a potential SNV. Thus, by missing such information, currently available databases and their applications in prioritizing and filtering variants for association analyses especially those with a large number of candidate SNVs are limited. Lastly, none of these databases included tissue specific information. It has been well studied that miRNAs have differential expression levels across different tissues (Ludwig et al., 2016). Thus, the functionalities of miRNA and MTS SNVs are greatly associated with the environment being considered.

To bridge these gaps, we have established a comprehensive database with all putative SNVs that might have an influence on miRNA targeting. We first compiled a collection of all possible SNVs in the 3’UTR of mRNAs that may disrupt a MTS or gain a new MTS based on predictions from three popular miRNA target prediction tools, namely TargetScan (Agarwal et al., 2015, http://www.targetscan.org/vert_70/, v7.0), miRanda (John et al., 2004, http://www.microrna.org/microrna/getDownloads.do, aug2010) and RNAhybrid (Rehmsmeier, Steffen, Höchsmann, Giegerich, & Ho, 2004, https://bibiserv.cebitec.uni-bielefeld.de/download/tools/rnahybrid.html, 2.1.1). At the same time, we calculated some miRNA-specific scores for all identified SNVs using these three miRNA target prediction tools. We next collected their corresponding prediction scores from multiple popular SNV functional annotation tools, such as CADD, DANN (Quang, Chen, & Xie, 2015), FATHMM-MKL (Shihab et al., 2015) and Eigen. Lastly, The Cancer Genome Atlas (TCGA) data were processed to obtain miRNA-mRNA correlation in multiple tissues for both normal and tumor samples. We named our database dbMTS (database of microRNA Target Site SNVs), which is the first known database that aims to include all putative SNVs in human 3’UTRs that may impact miRNA targeting along with their functional annotations. This database can help studies easily and quickly identify putative SNVs that may impact miRNA targeting and facilitate the prioritization of functional important SNVs in putative MTS at genome level. We have provided dbMTS as an attached database to dbNSFP which is available at: https://sites.google.com/site/jpopgen/dbNSFP.

## Data Sources and Processing

TargetScan v7.0, RNAhybrid, and miRanda were used to predict putative miRNA targets and to evaluate the effect of different SNVs on miRNA targeting. Briefly, these algorithms identify favorable miRNA binding sites by providing a numeric estimation of the likelihood and the binding efficacy for a specific miRNA-target pairing site. miRanda focuses more on the complementarity between the miRNA and the binding site. RNAhybrid focuses more on the minimum free energy hybridization between the miRNA and its target 3’UTR sequence. TargetScan adopts more comprehensive information from various aspects of the binding site: conservation of the target, context information such as the position of the site, and seed region complementarity etc. We chose these three algorithms for two reasons. First, they adopted different target prediction and scoring schemes, which enabled us to capture different aspects of miRNA targeting. Second, their executables were freely available online, so that we could make batch predictions locally.

The 3’-UTR coordinates and sequences were downloaded using the Table Browser utility from the UCSC genome browser. GENCODE (Harrow et al., 2012) gene annotation V23 basic set under genome assembly hg38 was retrieved, which included 3’-UTRs for 73,196 transcripts. miRNA sequence file for all species was downloaded from miRBase V21 at http://www.mirbase.org/ftp.shtml. Only human mature miRNAs were kept, which resulted in 2,588 mature miRNA sequences.

As the initial step to build dbMTS, our goal was to identify MTS SNVs and estimate their effect on miRNA targeting. To minimize the computational burden and at the same time capture the most impactful SNVs and their effect, we focused our research on those SNVs that pair with the miRNA seed region where a single mutation would completely disrupt the miRNA regulation. Our first step was to run the three miRNA target prediction algorithms with 2,588 human mature miRNAs and 3’UTR transcripts to get the reference miRNA targeting information in human (reference scores). Then, to estimate the SNVs’ effect on reference miRNA targetome, we would mutate each nucleotide of all the 3’UTR transcripts one-by-one and use these variant-induced 3’UTRs to run the three miRNA target prediction algorithms again (variant-induced scores; see **Supp. Figure S1** for detail). Next, we categorized all SNVs into three groups based on its estimated effect (**Figure 1**): 1) a SNV was classified as substitution when there are regulating miRNAs and have their seed regions overlap with this locus using both reference 3’UTR sequence and variant-induced 3’UTR sequence; 2) a SNV was classified as target loss where there are regulating miRNAs overlap with this locus using the reference 3’UTR sequence but not the variant-induced 3’UTR sequence; 3) a SNV was classified as target gain where there are regulating miRNAs overlap with this locus using the variant-induced 3’UTR sequence but not the reference 3’UTR sequence. For each SNV, the maximum difference between the reference score and variant-induced score was calculated to estimate how the miRNA targeting efficacy was changed after introducing the variant (**Figure 1**). Currently, there is no clear indication showing which of these three types of MTS SNVs is functionally more important. Thus, for each miRNA target prediction algorithm, we calculated rank scores within each type of the SNV to account for the possible impact of different scales of their raw scores between the three types of SNV groups.

**Figure 1.**
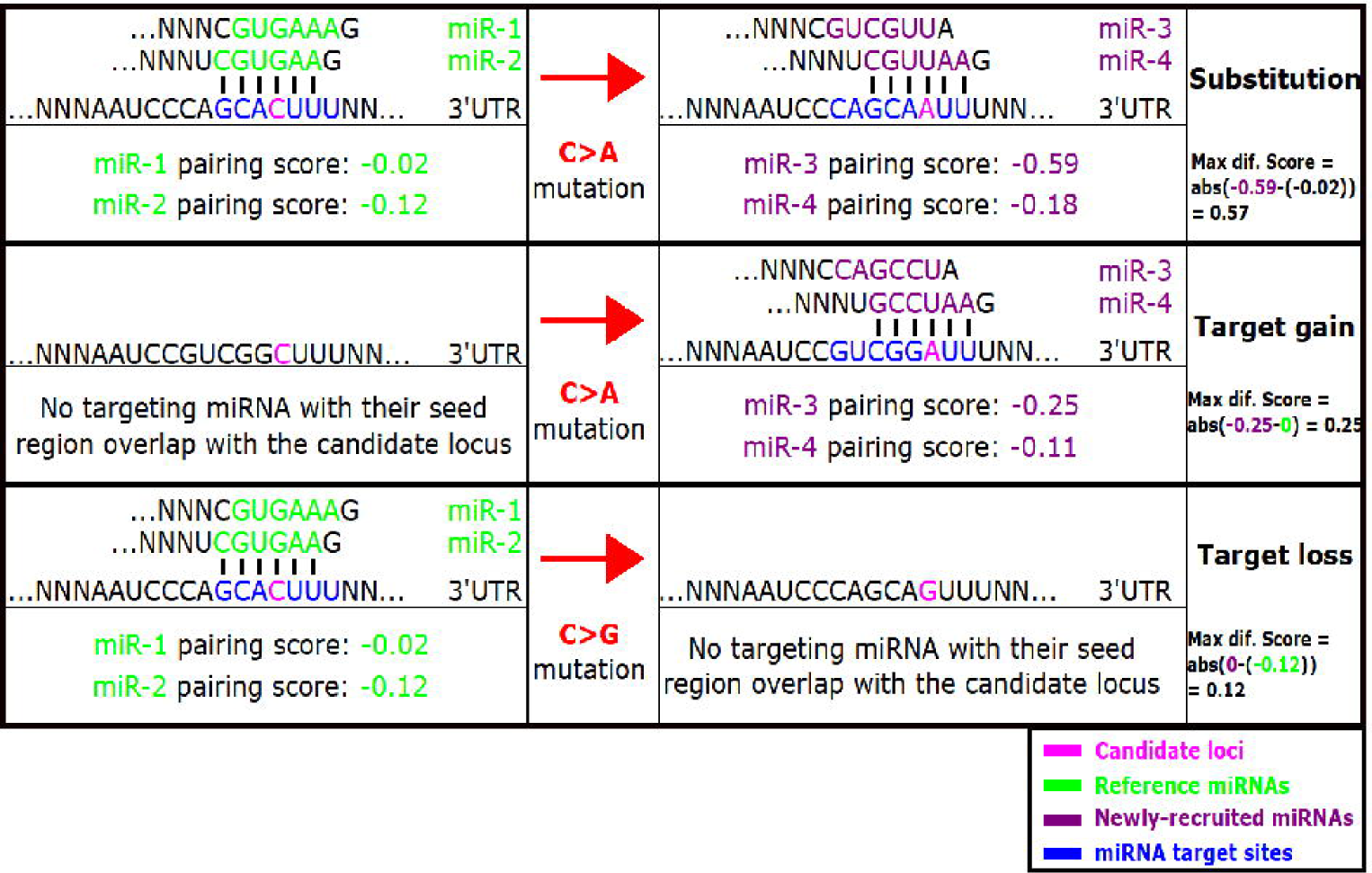
Illustration of the three types of SNVs and maximum difference score calculation using putative variants and putative miRNA and 3’UTR sequences. In first row, the C>A mutation caused substitution of targeting miRNAs. A substitution is when there were regulating miRNAs targeting with their seed regions overlapping with this locus, after introducing a mutation there was still at least one different miRNA targeting this locus. In second row, the C>A mutation caused target gain. A target gain is when there was no regulating miRNA, after introducing a mutation it gained at least one regulating miRNA. In third row, the C>G mutation caused target loss. A target loss is when there were regulating miRNAs, but after introducing a mutation there was no targeting miRNA. The maximum differences are calculated as abs(−0.59 – (−0.02)) = 0.57 for the C>A substitution SNV; abs(−0.25) = 0.25 for the C>A target gain SNV; abs(−0.12) = 0.12 for the C>G target loss SNV.

After identifying the three categories of SNVs that affect miRNA targeting for each of the three miRNA target prediction tools, these results were combined together to build the foundation of the database with all potential functional SNVs that could affect miRNA targeting. Then additional annotations were extract from Whole Genome Sequencing Annotator (WGSA) (X. Liu et al., 2015, https://sites.google.com/site/jpopgen/wgsa, v0.8) based on the positions of all the SNVs identified in our database. Some of the annotation categories include: functional consequences of genomic variants by VEP (McLaren et al., 2016); dbSNP variant IDs; GWAS Catalog entries; allele frequencies from various populations; clinical consequences from ClinVar; expression quantitative trait loci (eQTLs) from GTEx; mappability scores etc. In addition, major quantitative annotations which combined machine learning techniques with experimental information or other annotation scores were included. Moreover, for each miRNA-target pair, we have calculated the correlation of their expression within 15 different tissues from TCGA program. The expression data were obtained from (Bai et al., 2016) including 8 tissues from their published data and 7 tissues from their unpublished data. We provided these annotations to help users more easily rank and interpret a large number of candidate SNVs.

Among the large number of annotations included, in this study, we focused only on those popular quantitative measurements that had been proven to be useful under different scenarios to prioritize functional SNVs. However, please note that other annotations could potentially be as or more useful depending on specific research goals and interests. The 16 annotations we selected include eight conservation prediction scores: PhyloP46way primate, PhyloP100way vertebrate, PhyloP20way mammalian, PhastCons46way primate, PhastCons20way mammalian, PhastCons100way vertebrate, GERP_RS, and Siphy scores; eight integrative annotations that adopted more than one features or combined multiple individual annotations: integrated fitCons, FATHMM-MKL (coding and non-coding), Eigen, Eigen-PC, CADD, DANN and GenoCanyon scores.

## Database Contents

In our database, each SNV links to 221 unique fields with the first four columns as the primary identifier of the SNV: chromosome number, physical position on the chromosome as to hg38 (1-based coordinate), reference allele (as on the + strand), alternate allele (as on the + strand). For users who are interested, we also provided the identifier of the SNV as to hg19 human reference genome from column 5 to 8. Following the identifier information there are annotations we retrieved from WGSA (column 10 to 131). From column 132 to 221, there are exclusive information we obtained from the three miRNA target prediction tools and miRNA-target co-expression information. For each of the miRNA target prediction tools, there are 30 fields of information: 13 fields of predictions using reference 3’UTRs, 13 fields of predictions using SNV-induced 3’UTRs, and 4 fields with site level summary information: maximum difference score, its rank score, the transcript ID correspond to the maximum difference score, and the predicted category of the SNV. A more comprehensive description of these fields can be found at **Supp. Table S1**. For this study, detailed information of the 16 quantitative annotation scores and miRNA specific scores could be found at **Table 1**. The relatively low coverage of miRanda and RNAhybrid predictions resulted partly from their high threshold of reporting a ‘true’ MTS, and partly from built-in limitations of the program, e.g. RNAhybrid was not able to predict 3’UTR with length greater than 2000. Their results could be considered as a more constrained set of potential MTS SNVs.

**Table 1.** Annotations in dbMTS with their score range and missingness.

| **Annotations** | **min**<br>**score** | **max**<br>**score** | **No.**<br>**Missing** | **No.**<br>**SNVs** | **Percent non-**<br>**missing (%)** |
| --- | --- | --- | --- | --- | --- |
| phyloP46way_primate | -8.04 | 0.66 | 402463 | 131379720 | 99.69 |
| phyloP20way_mammalian | -12.51 | 1.2 | 161466 | 131620717 | 99.88 |
| phyloP100way_vertebrate | -20 | 10 | 153104 | 131629079 | 99.88 |
| phastCons46way_primate | 0 | 1 | 402460 | 131379723 | 99.69 |
| phastCons20way_mammalian | 0 | 1 | 161466 | 131620717 | 99.88 |
| phastCons100way_vertebrate | 0 | 1 | 153104 | 131629079 | 99.88 |
| GERP_RS | -12.3 | 6.17 | 2938733 | 128843450 | 97.77 |
| SiPhy_29way | 0 | 34.19 | 18703515 | 113078668 | 85.81 |
| integrated_fitCons | 0 | 0.84 | 151672 | 131630511 | 99.88 |
| GenoCanyon | 0 | 1 | 146368 | 131635815 | 99.89 |
| CADD | -6.41 | 35.5 | 596677 | 131185506 | 99.55 |
| DANN | 0.01 | 1 | 596677 | 131185506 | 99.55 |
| fathmm-MKL_non-coding | 0 | 1 | 633718 | 131148465 | 99.52 |
| fathmm-MKL_coding | 0 | 1 | 633535 | 131148648 | 99.52 |
| Eigen | -3.33 | 6.84 | 601953 | 131180230 | 99.54 |
| Eigen-PC | -3.38 | 17.69 | 601953 | 131180230 | 99.54 |
| miRanda_raw | 0 | 216 | 42475524 | 89306659 | 67.77 |
| miRanda_rankscore | 0 | 1 | 42475524 | 89306659 | 67.77 |
| TargetScan_raw | 0 | 11.07 | 1651761 | 130130422 | 98.75 |
| TargetScan_rankscore | 0 | 1 | 1651761 | 130130422 | 98.75 |
| RNAhybrid_raw | 0 | 60.8 | 74383715 | 57398468 | 43.56 |
| RNAhybrid_rankscore | 0 | 1 | 74383715 | 57398468 | 43.56 |

Correlation structures of the abovementioned quantitative annotation scores are shown in **Figure 2**. There was high correlation between conservation and most integrative scores, while there was little correlation between miRNA target prediction scores and all other scores. This could indicate that conservation or comparative genomic information was heavily used in these integrative algorithms, and those miRNA target prediction algorithms might be able to provide some additional information about functional importance of these SNVs regardless of conservation.

**Figure 2.**
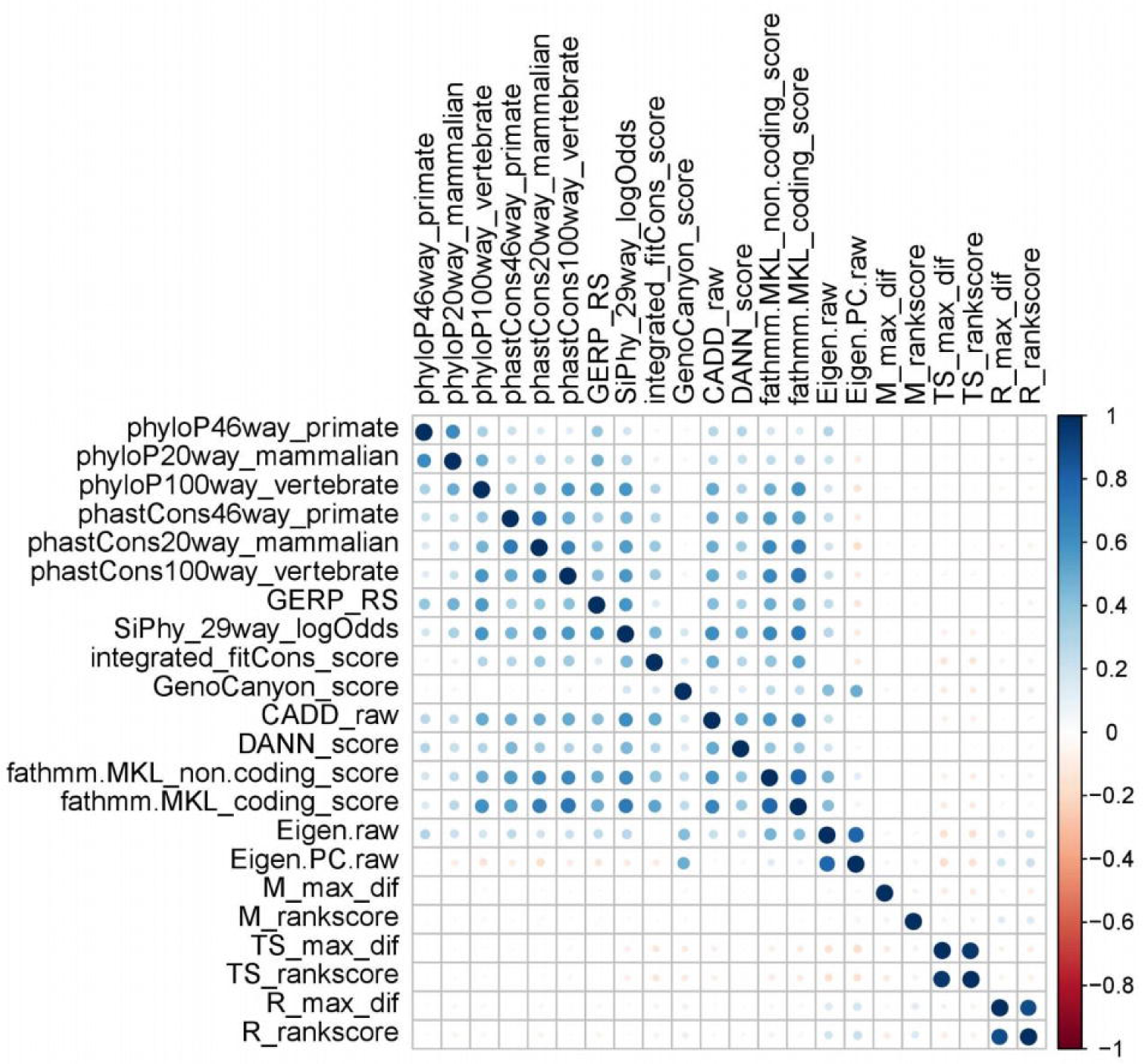
Correlation structure between different annotation scores.

## Utility and discussion

### Functional 3’UTR SNVs are more likely to fall in dbMTS

First, we checked if functionally important SNVs in 3’UTR are enriched in dbMTS. We retrieved two related datasets, namely ClinVar (Landrum et al., 2016, https://www.ncbi.nlm.nih.gov/clinvar/, 20160802) and the MiRNA SNP Disease Database or MSDD (Yue et al., 2018, http://www.bio-bigdata.com/msdd/home.jsp, June 2017). ClinVar is a public database with reported association between human variation and phenotypes. MSDD is a manually curated database containing experimentally supported associations between miRNA related SNVs and human diseases. For ClinVar, we identified 1,060 pathogenic SNVs in dbMTS, and an additional 1,797 SNVs in the rest of human 3’UTRs. Given that dbMTS includes 44,161,651 positions and human 3’UTRs have about 239,230,049 positions, it can be shown that pathogenic SNVs are over-represented in dbMTS (P < 0.00001). In addition, we extracted 118 unique SNVs in MSDD database that were labelled as 3’UTR variants. We found that seven of these SNVs were either located on chromosome X or resided in miRNA coding sequences which were not considered in our database. Among the remaining 111 SNVs, we were able to identify 109 of them in dbMTS (98.2%). Using the same 111 SNVs, PolymiRTS Database 3.0 and miRNASNP v2.0 covered 100 (90.1%) and 85 (77.3%) SNVs, respectively. These results indicated that dbMTS could identify potential SNVs that can affect miRNA targeting and are functionally important, which would provide users with increased ability to screen for functional SNVs in human 3’UTRs.

Second, we checked if our calculated miRNA specific scores (i.e. TS_rankscore, M_rankscore and R_rankscore) using the three miRNA prediction tools, TargetScan, miRanda and RNAhybrid, can further help user discriminate between non-functional and functional SNVs. Using MSDD, we found that SNVs with low TargetScan rank score (TS_rankscore < 0.2) showed statistically significant depletion (P < 0.05). This under-representation of low scores illustrated that when a SNV has a low TS_rankscore (<0.2), it is less likely to be functional, or at least less likely to be functionally enough to produce observable downstream mRNA expression change through impacting the miRNA targeting pathways.

### Comparison of predictive power between different annotation scores

After proving dbMTS’s ability to identify functional MTS SNVs, we next tried to compare some of the functional annotation scores in our database to check which one performs the best in separating potential functional MTS SNVs with non-functional ones. From ClinVar, we extracted 1,060 ‘pathogenic’ SNVs as a part of our true positive (TP) testing set and 2,939 ‘benign’ SNVs as our true negative (TN) testing set. From MSDD, we extracted the 109 unique SNVs that were labelled as 3’UTR variants and were identified in dbMTS. All SNVs identified at MSDD were labelled as TP in our testing dataset. Then testing samples extracted previously were combined. To ensure the SNVs being evaluated were completely non-coding and did not overlap with any coding regions, we removed those SNVs annotated as nonsynonymous or splicing by any of the three popular functional annotation tools, namely ANNOVAR (Wang, Li, & Hakonarson, 2010), VEP (McLaren et al., 2016) and SnpEff (Cingolani et al., 2012). Finally, we obtained a testing dataset with 160 TPs and 2,735 TNs. Using receiver operating characteristic (ROC) curves, we evaluated the performance of each annotation score in our database (**Figure 3**). To check if the imbalance testing set was an issue, we randomly selected 160 TNs and obtained similar result for each of the annotations. We found that the Eigen score had the overall best performance with the area under the curve (AUC) of 0.7335, followed by fathmm-MKL and several conservation scores. Interestingly, both TargetScan raw score and TargetScan rankscore had outperformed integrated fitCons score. This could probably be explained by the fact that currently available experimental data, such as those from the ENCODE project, was not comprehensive enough to correctly and fully capture miRNA binding information which highlighted the importance of such high-quality in-silico data (Consortium, 2012). For RNAhybrid and miRanda, all their predictions had AUCs with their 95% confidence interval including 0.5, meaning their predictive power to this testing dataset was no better than random guesses. This implied that relying solely on the difference between binding stabilities and the difference between base-pairing scores showed little predictive power for whether a MTS SNV was functional or not. Other context information around the binding site was also indispensable to predicting the efficacy of miRNA regulation and to infer SNV’s impact on miRNA targeting.

**Figure 3.**
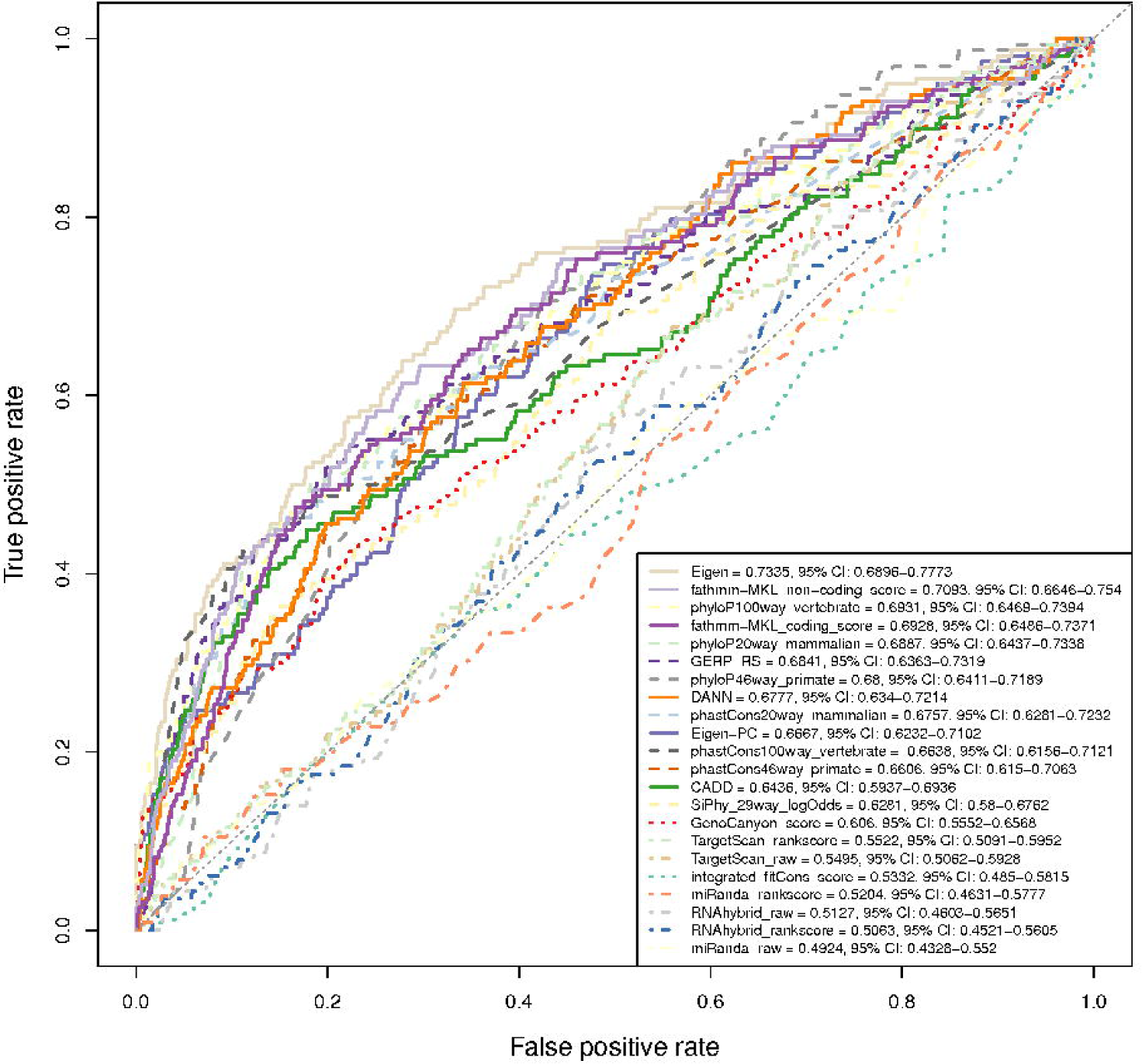
ROCs for functional annotation scores in our database using our curated testing set.

## Utilities and future studies

As mentioned previously, dbMTS included a large number of SNVs with their possible effects on miRNA targeting in the 3’UTR regions along with multiple functional annotation scores and predictions. Aside from simply using the overall best score, Eigen, another straightforward way to take advantage of the database is to use several annotation scores at once to find consensus predictions among them. This can be applied in two ways: the first way is to find consensus SNVs predicted by multiple miRNA target prediction algorithms to identify a stringent subset of SNVs that affect miRNA targeting; another way is to prioritize functional SNVs by using the predicted functional importance from multiple annotations (a list of recommended cut-off points for some of the annotations can be found at **Supp. Table S2**). Using this method, studies interested in SNVs and MTS could filter out a large number of neutral SNVs and keep those highly confident SNVs that are more likely to affect miRNA targeting for further analyses. For example, we investigated variants reported in GWAS Catalog and found 1,127 3’UTR SNVs that could potentially affect miRNA targeting (**Supp. Table S3**), which are good starting points for future functional validation. Additionally, with miRNA-target co-expression information, we observed that 11,393,295 SNVs in our database are located in MTS of miRNA-target pairs which showed negative correlation in some tissues and no positive correlation in any of the 15 tissues being considered. Thus, these variants and their effect predicted by computational tools are more likely to be authentic. This co-expression information added another layer of evidence to help users screen for truly functional variants. Moreover, given the extensive involvement of miRNAs in oncogenesis (Esquela-Kerscher & Slack, 2006; Lin & Gregory, 2015), using this same approach our database can be used to prioritize candidate driver mutations in cancer genomes. The richness of the available information can easily be used to further boost the user’s power to interpret non-coding SNVs. For example, eQTL loci can be used to associate SNVs and their targeting miRNAs with gene expression to gain a more well-rounded picture of gene regulation pathways.

Our database can be further improved in various ways. First, our database would benefit greatly from the future development of both miRNA target prediction tools and SNV functional annotation tools. Second, although we focused on SNVs, other types of genetic variations, such as insertions or deletions, can also disrupt miRNA targeting. Even though it is computationally expensive to evaluate their effects, including these types of mutations can undoubtedly further increase the comprehensiveness of our database. Third, since our database contained miRNA-specific raw scores from the three miRNA target prediction tools, they could be used to construct new measurements of functional importance other than the maximum potential difference we used in our study. Currently, our database is freely available at the dbNSFP website (https://sites.google.com/site/jpopgen/dbNSFP). We are planning to add a web portal which will enable users to search for the entries in the database using any of the following fields: genomic position, mature miRNA name or Ensembl transcript ID.

## Conclusion

In this study, we took advantage of three miRNA target prediction tools (TargetScan, miRanda and RNAhybrid) to identify all possible SNVs that could affect miRNA targeting in the 3’UTR of human mRNAs. We calculated the functional importance using the three above-mentioned tools and collected multiple popular functional annotation scores for these SNVs. In addition, we compared these functional annotation scores collected regarding their performance using a combined testing dataset. We found that Eigen outperformed all other individual annotations, and TargetScan showed statistically significant (though weak) predictive power regarding SNVs’ pathogenicity. We hope the presented database could facilitate researches interested in using MTS to prioritize functional SNVs or interpret of WGS results.

