## Supplemental material for "dbMTS: a comprehensive database of putative human microRNA target site SNVs and their functional predictions"

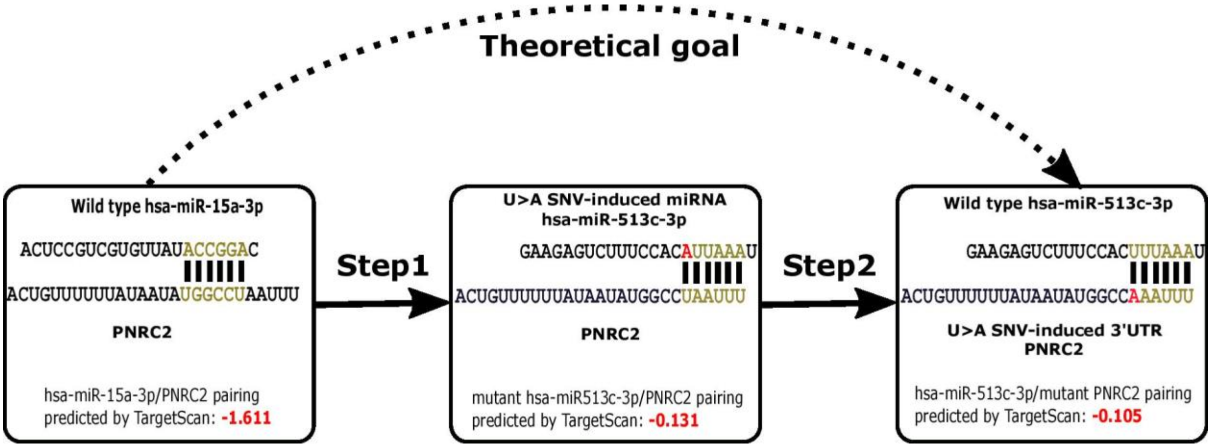


**Supp. Figure S1**. SNV in our database located at chr1:23962771. Given a locus of interest (U), our theoretical goal is to mutate U>A on the 3'UTR of gene PNRC2 (step 2). Given the seed region complementarity, it is much more computationally efficient to mutate U>A on has-miR-514-3p (step 1). As was shown from the illustration, step1 and step2 have almost identical pairing and have similar predicted scores. Thus, in this study, we used the method in step 1 to estimate the variant introduced in step 2.

**Supp. Table S1.** Description of 221 fields in dbMTS

Large table. See separate file.

**Supp. Table S2.** Recommended thresholds for 17 functional annotation scores in dbMTS

| **Annotation Scores** | **Thresholds** |
|---|---|
| phyloP46way_primate | 0.3755 |
| phyloP20way_mammalian | 0.613 |
| phyloP100way_vertebrate | 0.7275 |
| phastCons46way_primate | 0.1325 |
| phastCons20way_mammalian | 0.7405 |
| phastCons100way_vertebrate | 0.8505 |
| GERP_RS | 2.175 |
| SiPhy_29way_logOdds | 8.3847 |
| integrated_fitCons_score | 0.1480705 |
| GenoCanyon_score | 0.9999728 |
| CADD | 0.7570625 |
| DANN | 0.7046078 |
| fathmm-MKL_non-coding_score | 0.216635 |
| fathmm-MKL_coding_score | 0.1744 |
| Eigen | 0.2936396 |
| Eigen-PC | -0.09158359 |
| TS_rankscore | 0.5375 |

**Supp. Table S3.** 1,127 SNVs reported in GWAS Catalog that could potentially affect miRNA targeting

Large table. See separate file.
